# Cellpose: a generalist algorithm for cellular segmentation

**DOI:** 10.1101/2020.02.02.931238

**Authors:** Carsen Stringer, Tim Wang, Michalis Michaelos, Marius Pachitariu

## Abstract

Many biological applications require the segmentation of cell bodies, membranes and nuclei from microscopy images. Deep learning has enabled great progress on this problem, but current methods are specialized for images that have large training datasets. Here we introduce a generalist, deep learning-based segmentation method called Cellpose, which can precisely segment cells from a wide range of image types and does not require model retraining or parameter adjustments. We trained Cellpose on a new dataset of highly-varied images of cells, containing over 70,000 segmented objects. We also demonstrate a 3D extension of Cellpose which reuses the 2D model and does not require 3D-labelled data. To support community contributions to the training data, we developed software for manual labelling and for curation of the automated results, with optional direct upload to our data repository. Periodically retraining the model on the community-contributed data will ensure that Cellpose improves constantly.

## Introduction

Quantitative cell biology requires measurements of multiple cellular properties such as shape, position, RNA expression and protein expression [1]. To assign these properties to individual cells, one must first segment an image volume into cell bodies, usually based on a cytoplasmic or membrane marker [2–8]. This step can be straightforward when cells are sufficiently separated from one other, e.g. in monodispersed cultures in vitro. However, in many tissues, cells are tightly packed together and difficult to separate computationally.

Most methods for cell body segmentation make trade-offs between flexibility and automation. These methods range from fully-manual labelling [9], to user-customized pipelines involving a sequence of image transformations with user-defined parameters [2, 8, 10, 11], to fully automated methods based on deep neural networks with parameters estimated on large training datasets [4, 5, 7, 12, 13]. Fully automated methods have many advantages, such as reduced human effort, increased reproducibility and better scalability to big datasets from large screens. However, these methods are typically trained on specialized datasets, and do not generalize well to other types of data, requiring new human-labelled images to achieve best performance on any one image type.

Recognizing this generalization problem in the context of nuclear segmentation, a recent Data Science Bowl challenge amassed a dataset of varied images of nuclei from many different laboratories [14]. Methods trained on this dataset can generalize much more widely than those trained on data from a single lab. The winning methods from the challenge were based on established computer vision algorithms like the Mask R-CNN model [15, 16]. Following the competition, the dataset generated further progress, with other methods like Stardist and nucleAIzer being developed [17, 18].

In this work, we followed the approach of the Data Science Bowl team to collect and segment a large dataset of cell images from a variety of microscopy modalities and fluorescent markers. We had no guarantee that we could replicate the success of the nuclear dataset, because cells have highly diverse shapes and a single model may not be able to segment all these shapes. Previous approaches did not perform well on this dataset. We hypothesized that their failure was due to low expressive power, i.e. insufficient flexibility in describing diverse cell shapes. We therefore developed a new model with better expressive power called Cellpose. We next describe the architecture of Cellpose and the new training dataset, and then proceed to show performance benchmarks on test data, as well as an extension to 3D. The Cellpose package with a graphical user interface (Figure S1) can be installed from www.github.com/mouseland/cellpose or tested online at www.cellpose.org.

## Results

### Model design

In classical segmentation approaches like the watershed [19], the grayscale values of an image create a topological map, whose basins represent the segmented regions. These methods work well when the segmented objects are blob-like, so that they form smooth basins. However, many types of cells are not blob-like, which motivated us to construct an intermediate image representation that forms a smooth topological map (Figure 1ab).

**Figure 1:**
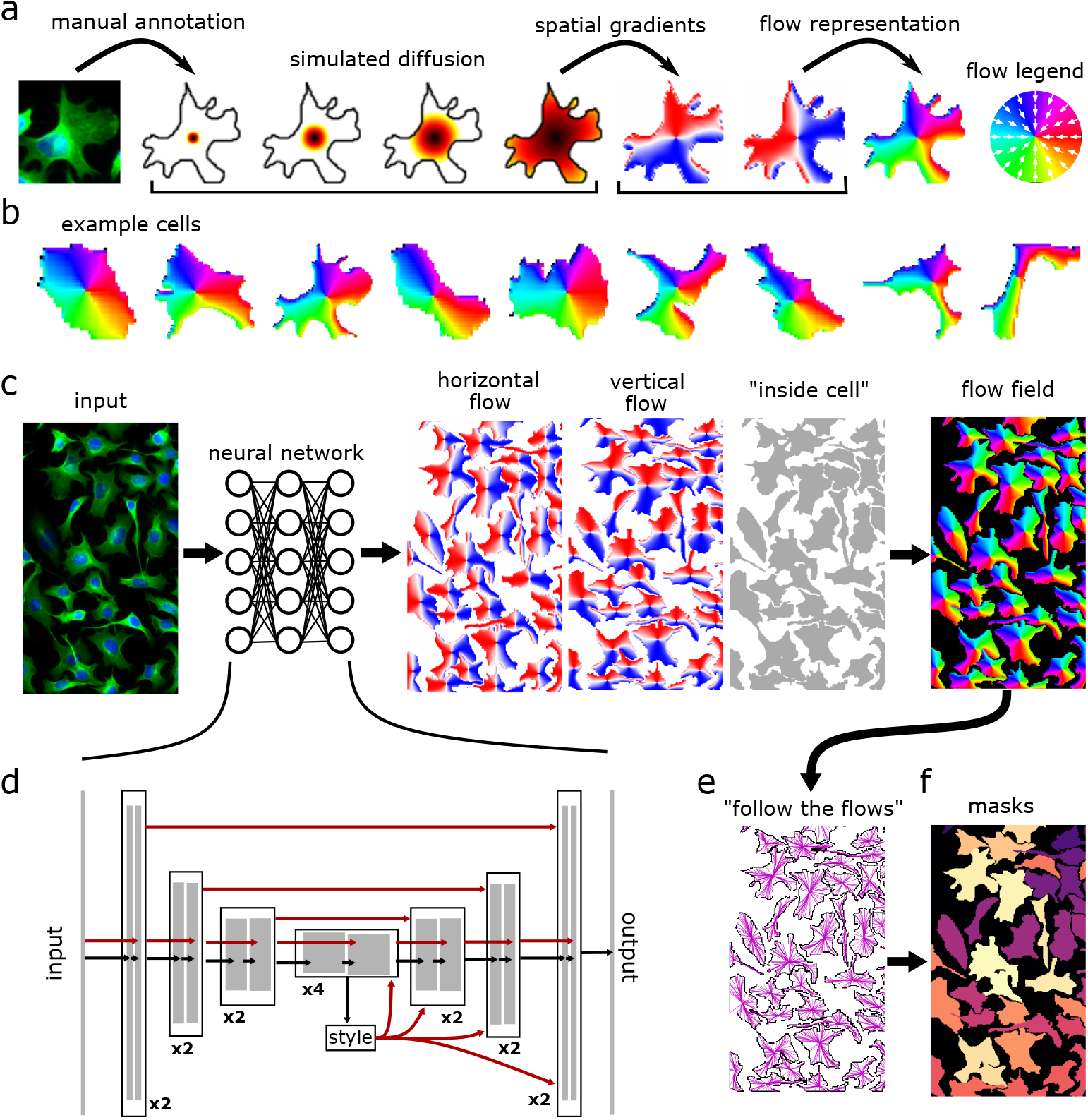
Model architecture. **a**, Procedure for transforming manually annotated masks into a vector flow representation that can be predicted by a neural network. A simulated diffusion process started at the center of the mask is used to derive spatial gradients that point towards the center of the cell, potentially indirectly around corners. The X and Y gradients are combined into a single normalized direction from 0° to 360°. **b,** Example spatial flows for cells from the training dataset. **cd,** A neural network is trained to predict the horizontal and vertical flows, as well as whether a pixel belongs to any cell. The three predicted maps are combined into a flow field. **d** shows the details of the neural network which contains a standard backbone neural network that downsamples and then upsamples the feature maps, contains skip connections between layers of the same size, and global skip connections from the image styles, computed at the lowest resolution, to all the successive computations. **e,** At test time, the predicted flow fields are used to construct a dynamical system with fixed points whose basins of attraction represent the predicted masks. Informally, every pixel “follows the flows” along the predicted flow fields towards their eventual fixed point. **f,** All the pixels that converge to the same fixed point are assigned to the same mask.

Our topological maps were generated from ground truth masks, manually drawn by a human annotator, through a process of simulated diffusion (Figure 1a). A neural network was then trained to predict (1) the horizontal gradients of the topological maps, (2) the vertical gradients, and finally (3) a probability map which indicates if a given pixel is part of any cells (Figure 1 c,d). On test images, the neural network predicted the horizontal and vertical gradients, which form vector fields or “paths”. By following these paths, all pixels belonging to a given cell should be routed to its center. Thus, by grouping pixels that flowed to the same center point, we could simultaneously segment individual cells and recover their shapes (Figure 1e,f, and Methods). The cell shapes were further refined by removing pixels with cell probabilities less than 0.5.

The neural network that predicts the spatial flows was based on the general U-net architecture, which downsamples the convolutional maps several times before upsampling in a mirror-symmetric fashion (Figure 1d). On the upsampling pass, the U-net “mixes in” the convolutional maps from the downsampling pass. This mixing is typically done by feature concatenation, but we opted for direct summation in order to reduce the number of parameters. We also replaced the standard building blocks of a U-net with residual blocks, which have been shown to perform better, and doubled the depth of the network as typically done for residual networks [20]. In addition, we used global average pooling on the smallest convolutional maps to obtain a representation of the “style” of the image (for similar definitions of style see [18, 21, 22]). We anticipated that images with different styles might need to be processed differently, and therefore fed the style vectors into all the following processing stages to potentially re-route and re-adjust the computations on an image by image basis.

We also use several test time enhancements to further increase the predictive power of the model: test time resizing, ROI quality estimation, model ensembling, image tiling and image augmentation (see Methods).

### Training dataset

We collected images from a variety of sources, primarily via internet searches for keywords such as “cyto-plasm”, ‘‘cellular microscopy”, ‘‘fluorescent cells” etc. Some of the websites with hits shared the images as educational material, promotional material or news stories, while others contained datasets used in previous scientific publications (i.e. OMERO [23, 24]). The dataset consisted primarily consisted of fluorescently labelled proteins that localized to the cytoplasm with or without DAPI-stained nuclei in a separate channel (n = 316 images). We also included images of cells from brightfield microscopy (n = 50) and membrane-labelled cells (n = 58). Finally, we included a small set of images from other types of microscopy (n = 86), and a small set of non-microscopy images which contained large numbers of repeated objects such as fruits, rocks and jellyfish (n = 98). We hypothesized that the inclusion of such images in the training set would allow the network to generalize more widely and more robustly.

We visualized the structure of this dataset by applying t-SNE to the image styles learned by the neural network (Figure 2) [25, 26]. Images were generally placed in the embedding according to the categories defined above, with the non-cell image categories scattered across the entire embedding space. To manually segment these images, we developed a graphical user interface in Python (Figure S1) that relies on PyQt and the pyqtgraph package [27, 28]. The interface enables quick switching between views such as outlines, filled masks and image color channels, as well as easy image navigation like zooming and panning. A new mask is initiated with a right-click, and gets completed automatically when the user returns within a few pixels of the starting area. This interface allowed human operators to segment 300-600 objects per hour, and is part of the code we are releasing with the Cellpose package. The automated algorithm can also be run in the same interface, allowing users to easily curate and contribute their own data to our training dataset.

**Figure 2:**
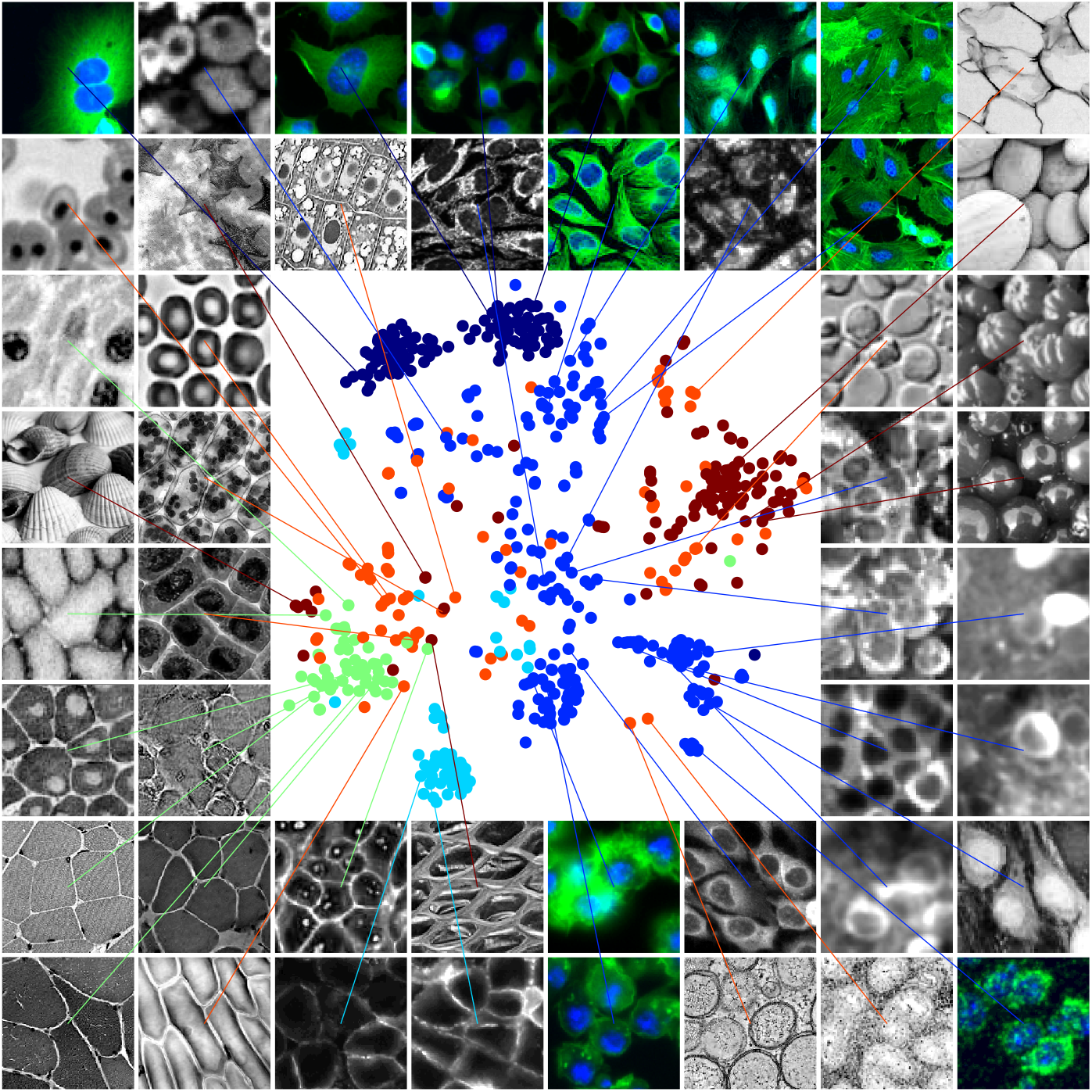
Visualization of the diverse training dataset. The styles from the network for all the images in the cytoplasm dataset were embedded using t-SNE. Each point represents a different image. Legend: dark blue = CellImageLibrary, blue = cytoplasm, cyan = membrane, green = non-fluorescent cells, orange = microscopy other, red = non-microscopy. Randomly chosen example images are shown around the t-SNE plot. Images with a second channel marking the nucleus are displayed in green/blue.

Within the varied dataset of 616 images, a subset of 100 images were pre-segmented as part of the CellImageLibrary [29]. These two-channel images (cytoplasm and nucleus) were from a single experimental preparation and contained cells with complex shapes. We reasoned that the segmentation methods we tested would not overfit on such a large and visually uniform dataset, and may instead fail due to a lack of expressive power such as an inability to precisely follow cell contours, or to incorporate information from the nuclear channel. We therefore chose to also train the models on this dataset alone, as a “specialist” benchmark of expressive power (Figure 3a).

**Figure 3:**
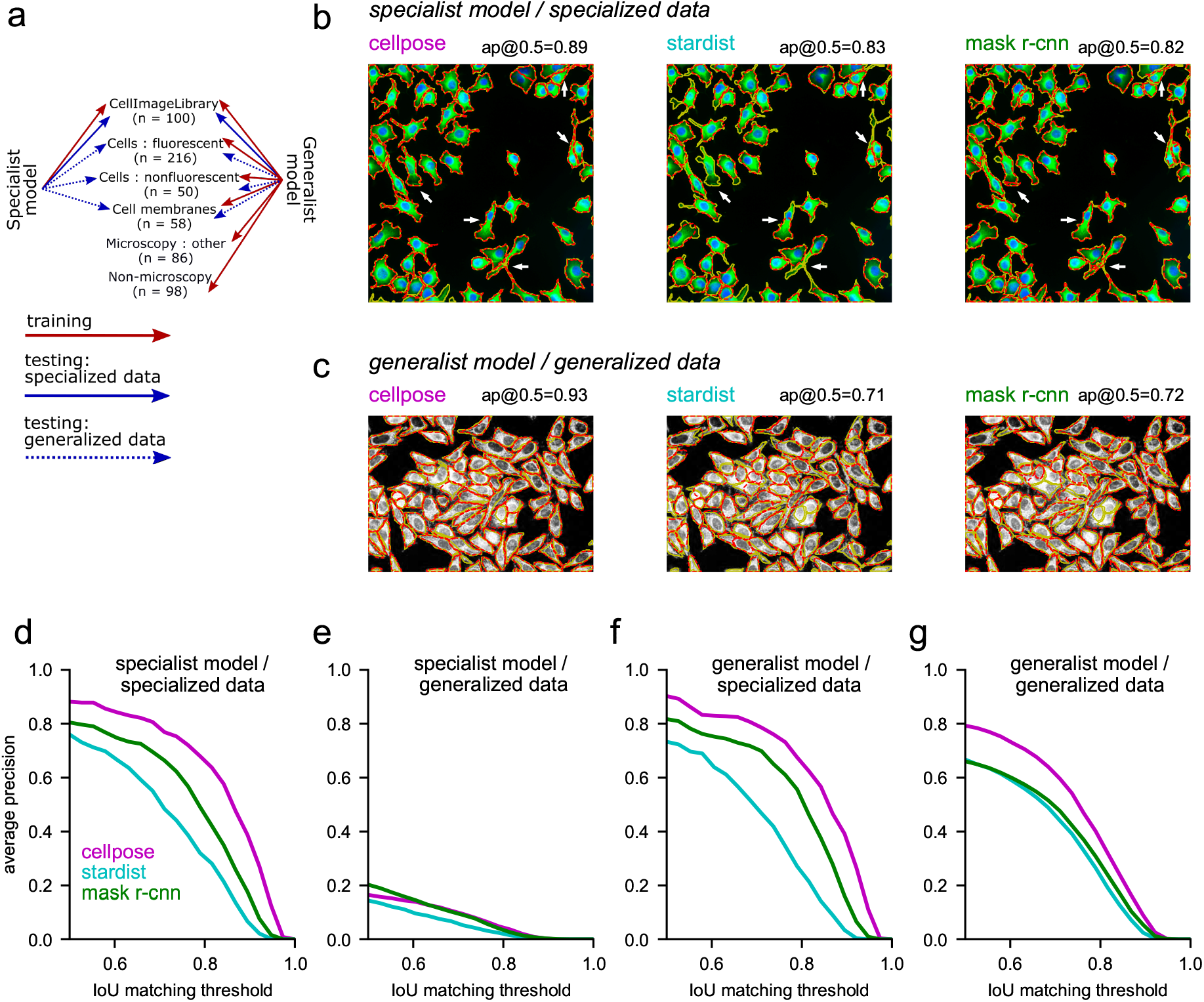
Segmentation performance of specialist and generalist algorithms. **a**, Illustration of the training and test data for specialist and generalist algorithms. We refer to the CellImageLibrary dataset as a ‘specialist dataset” due to the uniformity of its 100 images. **b,** Example test image segmentation for Cellpose, Stardist and Mask R-CNN, when trained as specialist models. **c,** Example test image segmentation for the same models trained as generalist models. **defg,** Quantified performance of all models, trained as specialists or generalists, and tested on specialized or generalized data. The IoU threshold (intersection over union) quantifies the match between a predicted mask and a ground truth mask, with 1 indicating a pixel-perfect match and 0.5 indicating as many correctly matched pixels as there were missed and false positive pixels. The average precision score is computed from the proportion of matched and missed masks. To gain an intuition for the range of these scores, refer to the reported values in **bc** as well as Figure 4. **e,** Note the poor generalization of the specialist models. **f,** Note the similar performance of the generalist and specialist model on the specialist data (compare to d). **g,** The generalist cellpose model outperforms the other models by a large margin on generalized data.

### Benchmarks

For the specialist benchmarks, we trained Cellpose and two previous state of the art methods, Stardist [17] and Mask-RCNN [15, 16], on the CellImageLibrary (Figure 3b). On test images, we matched the predictions of the algorithm to the true masks at different thresholds of matching precision, defined as the standard intersection over union metric (IoU). We evaluated performance with the standard average precision metric (*AP*), computed from the numbers of true positives *TP*, false positives *FP* and false negatives *FN* as 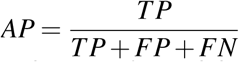. We found that Cellpose significantly outperformed the previous methods at all matching thresholds. For example, at the commonly used IoU threshold of 0.5, Cellpose correctly matched 485 out of 521 total ground truth ROIs, and gave only 18 false positives. This corresponded to an average precision of 0.88, compared to 0.76 for Stardist and 0.80 for Mask R-CNN. At higher IoUs, which benchmark the ability to precisely follow cell contours, the relative improvement of Cellpose compared to the other methods grew even larger. This analysis thus shows that Cellpose has enough expressive power to capture complicated shapes (Figure 3d).

However, the models trained on the specialized data performed poorly on the full dataset (Figure 3e), motivating the need for a generalist algorithm. To help with generalization across different image resolutions, we resized all images in the training set to the same average cell size. To estimate cell size on test images, we trained another model based on the image style vectors (see methods, Figure S2). Users may also manually input the cell size to the algorithm.

Across all image types (specialized and generalized cell data), we found that the generalist Cellpose model had an average precision of 0.82 at a threshold of 0.5, significantly outperforming Stardist and Mask R-CNN which had average precisions of 0.68 and 0.70 respectively. Example segmentations are shown for one image in Figure 3c and for 36 other images in Figure 4 for Cellpose, Figure S3 for Stardist and Figure S4 for Mask R-CNN. On the specialized data alone (the CellImageLibrary), the generalist Cellpose model matched the performance of the specialist model (Figure 3f). This shows that the inclusion of the other images in the training set did not saturate the capacity of the network, implying that Cellpose has spare capacity for more training data, which we hope to identify via community contributions. For the generalized data alone (all the cell images except those in the CellImageLibrary), Cellpose had an average precision of 0.79, while Stardist and Mask-RCNN had average precisions of 0.67 and 0.66 respectively (IoU threshold = 0.5, Figure 3g). All models had relatively worse performance on “non-microscopy” and “microscopy:other” images, with Cellpose scoring 0.73 compared to 0.54 and 0.55 for Stardist and Mask R-CNN (IoU threshold = 0.5, Figure S5). These relatively worse scores are likely due to images in these categories being highly visually distinct from other images in the training set. Note that the advantage of Cellpose compared to other models grew on these visually-distinct images, suggesting that Cellpose has better generalization performance.

**Figure 4:**
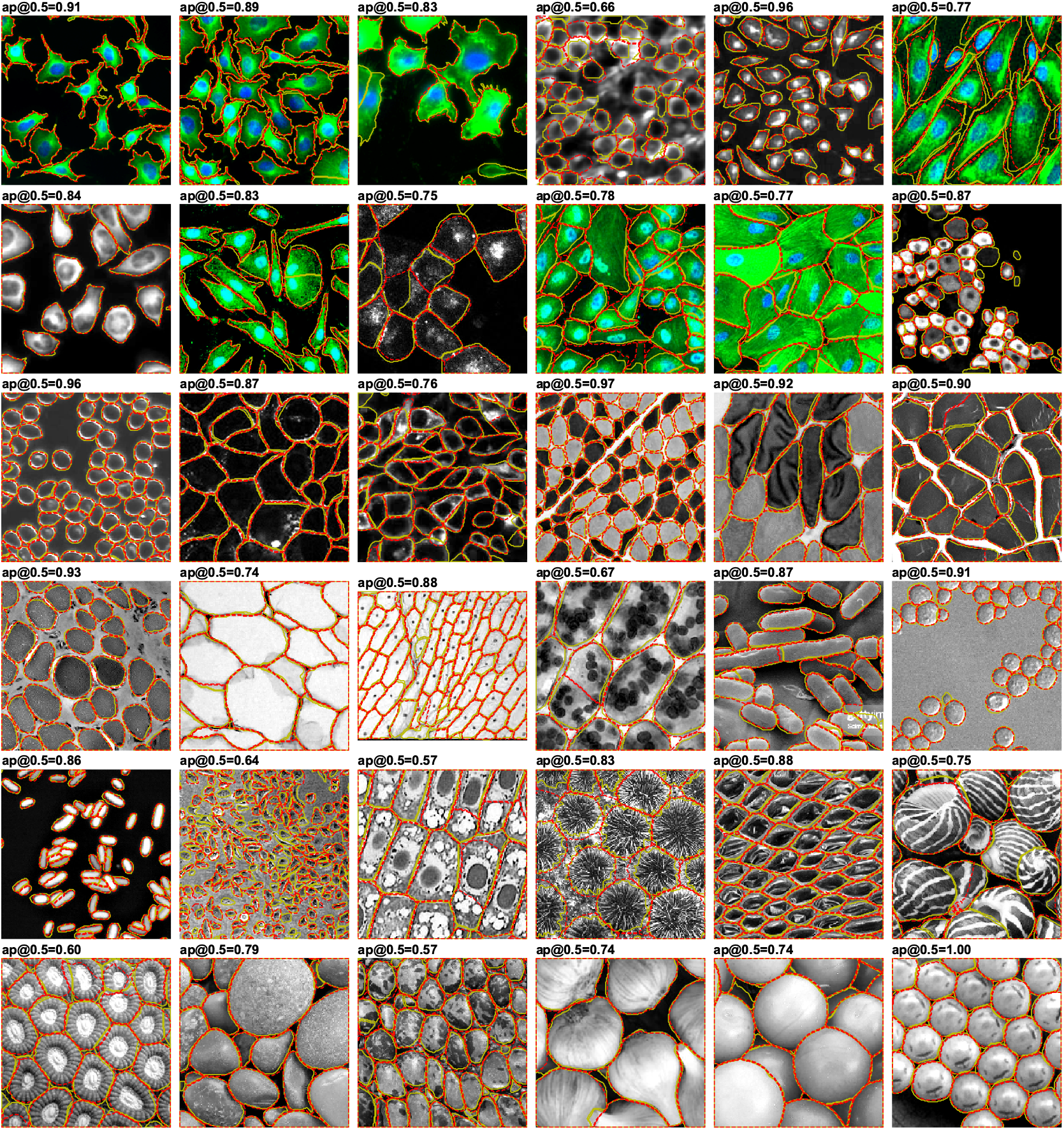
Example cellpose segmentations for 36 test images. The ground truth masks segmented by a human operator are shown in yellow, and the predicted masks are shown in dotted red line. Compare to Stardist and Mask-RCNN in Figure S3 and Figure S4.

Finally, we assembled a large dataset of images of nuclei, by combining pre-segmented datasets from several previous studies, including the Data Science Bowl kaggle competition [14]. Because nuclear shapes are simpler, this dataset did not have as much variability as the dataset of cells, as illustrated by the t-SNE embedding of the segmentation styles (Figure S6a). Cellpose outperformed the other methods on this dataset by smaller amounts, likely reflecting the simpler shapes of the masks, and the more uniform aspect of the nuclei (Figure S6b-d, Figure S7).

### 3D segmentation

Our last contribution is to generalize Cellpose to threedimensional segmentation. This task usually requires 3D training data, which is much more difficult to obtain than 2D training data, as it consists of one 2D segmentation for every Z position in the volume. At the median cell diameter size of 30 pixels in our dataset, we can estimate that every cell would take 30 times longer to manually segment in 3D than in a single 2D slice.

We designed a new method for extending Cellpose to 3D, using only the trained 2D model and no additional 3D training data. For a test volume, we ran Cellpose on all XY, XZ and YZ slices independently (Figure 5ab). For each point, this procedure generated two estimates of the gradient in X, Y and Z (i.e. 6 total predictions), which we then averaged together to obtain a complete set of 3D vector flows. To generate ROIs, we proceed like before to run the pixel dynamics and evaluate which pixels converge together to the same fixed points. Those pixels are then assigned to the same mask (Figure 5e). For comparison, we used ilastik to generate a 3D segmentation pipeline specific for this volume. The parameters of ilastik were chosen manually to give good performance on a subset of the volume that was not used for testing.

**Figure 5:**
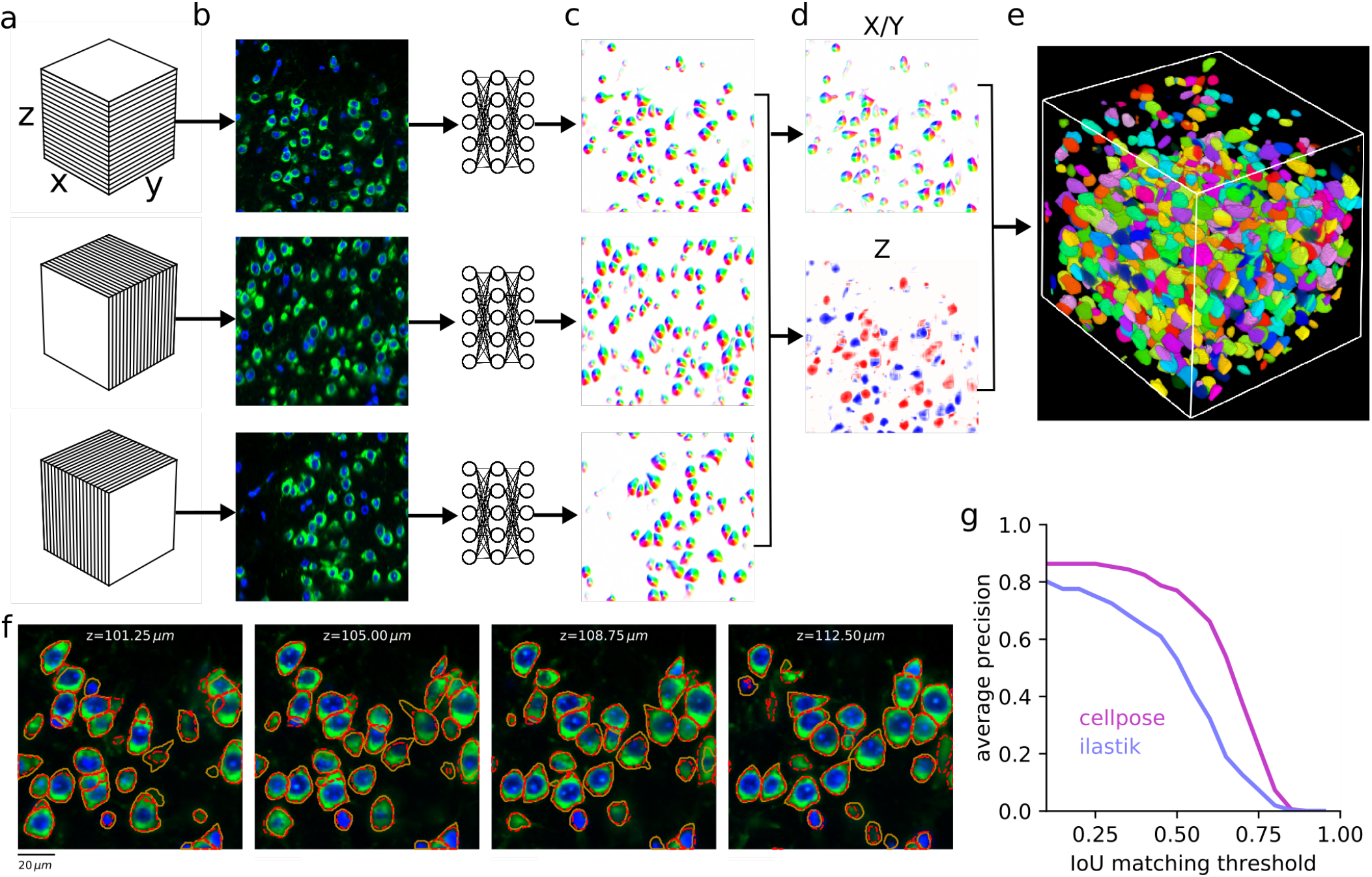
Segmentation in 3D without 3D labelled data. **a**, A 3D volume is sliced into XY, XZ and YZ sections respectively, **b,** Example frames for each type of section. **c,** Each 2D frame is passed through the cellpose prediction network, generating predicted 2D flows. **d,** The six predicted flow maps (two each for X, Y and Z) are pairwise averaged and combined into a single 3D flow field XYZ. Shown is the XY flow map, and the Z flow map for an XY slice through the volume. **e,** The 3D flows are used to create pixel dynamics towards the 3D sinks of the flow field, and the pixels are grouped into 3D masks if they converge to the same sink. **f,** Example XY slices with ground truth (yellow) and predicted (red) masks at different depths in the Z-stack. **g,** Matching performance at different IoU thresholds.

We evaluated the performance of Cellpose by comparing to manual annotations of a test 3D volume in which the DNA and RNA were co-stained to serve as nuclear and cytoplasmic markers, respectively (Figure 5f). The human annotator found 183 cells, while Cellpose predicted 172 cells, with 17 false positives at an IoU threshold of 0.5 (Figure 5g, Movie 1,2). At all IoU thresholds, the generalist Cellpose 3D model outperformed the ilastik pipeline which was manually optimized for this dataset.

## Discussion

Here we introduced Cellpose, a generalist model that can segment many types of cells, without requiring parameter adjustments, new training data or further model retraining. Cellpose uses two major innovations: a reversible transformation from training set masks to vector flows that can be predicted by a neural network, and a large segmented dataset of varied images of cells. In addition, multiple smaller improvements to the basic approach led to a significant cumulative performance increase: we developed methods to use an image “style” for changing the neural network computation on an image by image basis, to validate segmented ROIs, to average the predictions of multiple models, to resize images to a common object size, to average model predictions in overlapping tiles, and to augment those predictions by flipping the tiles horizontally and vertically. Finally, we introduced a new method for 3D cell segmentation that reuses the 2D model and does not require new 3D labelled data.

Our approach to the segmentation problem allows anyone to contribute new training data to improve Cellpose for themselves and for others. We encourage users to contribute up to a few manually segmented images of the same type, which we will use to periodically retrain a single generalist Cellpose model for the community. Cellpose has high expressive power and high capacity, as shown by its ability to segment cells with complex protuberances like elongated dendrites, and even non-cell objects like rocks and jellyfish. We therefore expect Cellpose to successfully incorporate new training data that has a passing similarity to an image of cells and consists of collections of objects that resemble each other. In contrast, we do not expect Cellpose to work well on images where different objects have different shapes and textures or when occlusions require highly overlapping masks.

Other extensions of Cellpose are possible, along the same principles as the 3D extension. One such application might be cell tracking, which could be addressed by adding a ‘‘temporal flow” dimension to the spatial flow dimensions. Combined with new histology methods like spatial transcriptomics [30], Cellpose has the potential to aid and enable novel approaches in quantitative single-cell biology that are reproducible and scalable to large datasets.

## Acknowledgments

This research was funded by the Howard Hughes Medical Institute at the Janelia Research Campus. We thank Paul Tillberg and members of the Tillberg lab for advice related to expansion microscopy, and Weinan Sun for sharing calcium imaging data from mouse hippocampus. We also thank Stephan Saalfeld and Jan Funke for useful discussions.

## Data and code availability

The code and GUI are available at www.github.com/mouseland/cellpose. To test out the model directly on the web, visit www.cellpose.org. Note that the test time augmentations and tiling, which improve segmentation, are not performed on the website to save computation time. The manually segmented cytoplasmic dataset will be made available upon publication, and will be continuously updated with community-contributed data.

## Methods

The Cellpose code library is implemented in Python 3 [31], using numpy, scipy, numba, opencv and the deep learning package mxnet [32–36]. The graphical user interface additionally uses PyQt, pyqtgraph and scikit-image [27, 28, 37]. The benchmark values were computed using the stardist implementation of average precision [38]. The figures were made using matplotlib and jupyter-notebook [39, 40].

### Datasets

#### Specialized data

This dataset consisted of 100 fluorescent images of cultured neurons with cytoplasmic and nuclear stains obtained from the CellImageLibrary [29]. Each image has a manual cytoplasmic segmentation. We corrected some of the manual segmentation errors where touching neurons were joined, and removed any overlaps in the segmentation so that each pixel in the image was assigned to at most one cytoplasmic mask. A small number of dendrites were cut off by this process, but the bulk of their cell bodies was un-modified.

#### Generalized data

The full dataset included the 100 images described above, as well as 516 additional images from various sources detailed below. All these extra images were fully manually segmented by a single human operator (MM). 69 of these images were reserved for testing.

216 of the images contained cells with fluorescent cytoplasmic markers. We used BBBC020 from the Broad Bioimage Benchmark Collection [41], which consisted of 25 images of bone-marrow derived macrophages from C57BL/6 mice. We used BBBC007v1 image set version 1 [42], which consisted of 15 fluorescent images of Drosophila melanogaster Kc167 cells with stains for DNA and actin. We imaged mouse cortical and hippocampal cells expressing GCaMP6 using a two-photon microscope, and included 8 images (mean activity) of such data. The other images in this set were obtained through Google image searches. We also used 10 images from confocal imaging of mouse cortical neurons with cytoplasmic and nuclear markers, from a similar dataset to that we used for 3D segmentation.

50 of the images were taken with standard bright-field microscopy. There were four images shared via OMERO [23] of pancreatic stem cells on a polystyrene substrate taken using a light microscope [43]. The other images in this set were obtained through google image searches.

In 58 of the images, the cell membrane was fluorescently labeled. We used the Micro-Net image set, which consisted of 40 fluorescent images of mouse pancreatic exocrine cells and endocrine cells with a membrane marker (Ecad-FITC) and a nuclear marker (DAPI) [44]. Because the nuclear marker did not appear in focus with the membrane labels, we discarded this channel, and re-segmented all images according to the membrane marker exclusively. The other images in this set were obtained through google image searches.

86 of the images were other types of microscopy samples which were either not cells, or cells with atypical appearance. These images were obtained through google image search.

98 of the images were non-microscopy images of repeating objects. These images were obtained through google image search, and include images of fruits, vegetables, artificial materials, fish and reptile scales, starfish, jellyfish, sea urchins, rocks, sea shells, etc.

#### Nucleus data

This data set consisted of 1139 images with manual segmentations from various sources, 111 of which were reserved for testing. We did not segment any of these images ourselves.

We used image set BBBC038v1 [14], available from the Broad Bioimage Benchmark Collection. For this image set, we used the unofficial corrected manual labels provided by Konstantin Lopuhin [45].

We used image set BBBC039v1 [14], which consists of 200 fluorescent images of U2OS osteosarcoma cells with the Hoechst stain. Some of these images overlap with the BBBC038v1, so we only used the 157 unique images.

We used image set MoNuSeg, which consists of 30 H&E stained histological images of various human organs [46]. Because the images contain many small nuclei (~700 per image), we divided each of these images into 9 separate images. We inverted the polarity of these images so that foreground nuclear pixels had higher intensity values than the background, which is more similar to fluorescence images.

We also used the image set from ISBI 2009, which consists of 97 fluorescent images of U2OS cells and NIH3T3 cells [47].

#### 3D data

We used a DNA stain (DAPI) and an RNA stain (fluorescent oligonucleotide probe against the 28S rRNA) to label the nucleus and cytoplasm respectively for a volume of murine cortical neurons. The tissue was first expanded by a factor of two and digested to remove all proteins [48, 49]. A small cube (~ 190×190×190 μm) was manually labelled by a human annotator in the Cellpose GUI. We allowed the annotator to potentially skip “easy” planes while drawing a cell, with an algorithm automatically interpolating the shapes of cells in those planes.

### Auxiliary vector flow representation

Central to the Cellpose model is an auxiliary representation which converts the cell masks to two images or “maps” of the same size as the original image. These maps are similar to the vector fields predicted by tracking models like OpenPose [50], which point from object parts to other object parts and are used in object tracking and coarse segmentation. In our case, the vector gradient flows lead all pixels within a cell towards its center. These flows do not necessarily point directly towards the center of the cell [51], because that direction might intersect cell boundaries. Instead, the flows are designed so that locally they translate pixels to other pixels inside cells, and globally over many iterations translate pixels to the eventual fixed points of the flows, chosen to be the cell centers. Since all pixels from the same cells end up at the same cell center, it is to recognize which pixels should be grouped together into ROIs, by assigning the same cell label to pixels with the same fates. The task of the neural network is to predict these gradients from the raw image, which is potentially a highly nonlinear transformation. Note the similarity to gradient flow tracking methods, where the gradients are computed directly as derivatives of the raw image brightness [52].

To create a flow field with the properties described above, we turned to a heat diffusion simulation. We define the “center” of each cell as the pixel in a cell that was closest to the median values of horizontal and vertical positions for pixels in that cell. Other definitions of “center”, such as the 2D medoid, resulted in similar-looking flows and algorithm performance. In the heat diffusion simulation, we introduce a heat source at the center pixel, which adds a constant value of 1 to that pixels value at each iteration. Every pixel inside the cell gets assigned the average value of pixels in a 3×3 square surrounding it, including itself, at every iteration, with pixels outside of a mask being assigned to 0 at every iteration. In other words, the boundaries of the cell mask are “leaky”. This process gets repeated for N iterations, where N is chosen for each mask as twice the sum of its horizontal and vertical range, to ensure that the heat dissipates to the furthest corners of the cell. The distribution of heat at the end of the simulation approaches the equilibrium distribution. We use this final distribution as an energy function, whose horizontal and vertical gradients represent the two flow fields that in our auxiliary vector flow representation.

### Deep neural network

The input to the neural network was a two-channel image with the primary channel corresponding to the cytoplasmic label, and the optional secondary channel corresponding to nuclei, which in all cases was a DAPI stain. When a second channel was not available, it was replaced with an image of zeros. Raw pixel intensities were scaled for each image so that the 1 and 99 percentiles corresponded to 0 and 1.

The neural network was composed of a downsampling pass followed by an upsampling pass, as typical in U-nets [3]. Both passes were composed of four spatial scales, each scale composed of two residual blocks, and each residual block composed of two convolutions with filter size 3×3, as is typical in residual networks [20]. This resulted in 4 convolutional maps per spatial scale, and we used max pooling to downsample the feature maps. Each convolutional map was preceded by a batchnorm + relu operation, in the order suggested by He et al, 2016 [53]. The skip connections were additive identity mappings for the second residual block at each spatial scale. For the first residual block we used 1×1 convolutions for the skip connections, as in the original residual networks [20], because these convolutions follow a downsampling/upsampling operation where the number of feature maps changes.

In-between the downsampling and upsampling we computed an image style [21], defined as the global average pool of each feature map [22], resulting in a 256-dimensional feature vector for each image. To account for differences in cell density across images, we normalized the feature vector to a norm of 1 for every image. This style vector was passed as input to the residual blocks on the upsampling pass, after projection to a suitable number of features equal to the number of convolutional feature maps of the corresponding residual block, as described below.

On the upsampling pass, we followed the typical U-net architecture, where the convolutional layers after an upsampling operation take as input not only the previous feature maps, but also the feature maps from the equivalent level in the downsampling pass [3]. We depart from the standard feature concatenation in U-nets and combine these feature maps additively on the second out of four convolutions per spatial scale [54]. The last three convolutions, but not the first one, also had the style vectors added, after a suitable linear projection to match the number of feature maps and a global broadcast to each position in the convolution maps. Finally, the last convolutional map on the upsampling pass was given as given as input to a 1×1 layer of three output convolutional maps. The first two of these were used to directly predict the horizontal and vertical gradients of the Cellpose flows, using an L2 loss. The third output map was passed through a sigmoid and used to predict the probability that a pixel is inside or outside of a cell with a cross-entropy loss. To match the relative contributions of the L2 loss and cross-entropy loss, we multiplied the Cellpose flows by a factor of 5.

We built and trained the deep neural network using mxnet [36].

#### Training

The networks were trained for 500 epochs with stochastic gradient descent with a learning rate of 0.2, a momentum of 0.9, batch size of 8 images and a weight decay of 0.00001. The learning rate was started at 0 and annealed linearly to 0.2 over the first 10 epochs to prevent initial instabilities. The value of the learning rate was chosen to minimize training set loss (we also tested 0.01, 0.05, 0.1 and 0.4). At epoch 400 the learning rate annealing schedule was started, reducing the learning rate by a factor of 2 every 10 epochs. For all analyses in the paper we used a base of 32 feature maps in the first layer, growing by a factor of two at each downsampling step and up to 256 feature maps in the layer with the lowest resolution. Separate experiments on a validation set held out from the main training dataset confirmed that increases to a base of 48 feature maps were not helpful, while decreases to 24 and 16 feature maps hurt the performance of the algorithm.

#### Image augmentations for training

On every epoch, the training set images are randomly transformed together with their associated vector fields and pixel inside/outside maps. For all algorithms, we used random rotations, random scaling and random translations, and then sampled a 224 by 224 image from the center of the resultant image. For Cellpose, the scaling was composed of a scaling to a common size of cells, followed by a random scaling factor between 0.75 and 1.25. For Mask R-CNN and Stardist, the common size resizing was turned off because these methods cannot resize their predictions. Correspondingly, we used a larger range of scale augmentations between 0.5 and 1.5. Note that Mask R-CNN additonally employs a multi-scale training paradigm based on the size of the objects in the image.

The random rotations were uniformly drawn from 0° to 360°. Random translations were limited along X and Y to a maximum amplitude of (*l_x_* – 224)/2 and (*l_y_* – 224)/2, where *l_y_* and *l_x_* are the sizes of the original image. This ensures that the sample is taken from inside the image. Rotations, but not translations and scalings, result in changes to the direction of vectors. We therefore rotated the mask flow vectors by the same angles the images were rotated.

#### Mask recovery from vector flows

The output of the neural network after tiling is a set of three maps: the horizontal gradients, the vertical gradients, and the pixel probabilities. The next step is to recover the masks from these. First, we threshold the pixel probability map and only consider pixels above a threshold of 0.5. For each of these pixels, we run a dynamical system starting at that pixel location and following the spatial derivatives specified by the horizontal and vertical gradient maps. We use finite differences with a step size of 1. Note that we do not re-normalize the predicted gradients, but the gradients in the training set have unit norm, so we expect the predicted gradients to be on the same scale. We run 200 iterations for each pixel, and at every iteration we take a step in the direction of the gradient at the nearest grid location. Following convergence, pixels can be easily clustered according to the pixel they end up at. For robustness, we also extend the clusters along regions of high-density of pixel convergence. For example, if a high-density peak occurs at position (*x,y*), we iteratively agglomerate neighboring positions which have at least 3 converged pixels until all the positions surrounding the agglomerated region have less than 3 pixels. This ensures success in some cases where the deep neural network is not sure about the exact center of a mask, and creates a region of very low gradient where it thinks the center should be.

### Test time enhancements

We use several test time enhancements to further increase the predictive power of the model: test time resizing, ROI quality estimation, model ensembling, image tiling and image augmentation. We describe them briefly in this paragraph, and in more detail below. We estimate the quality of each predicted ROI according to the discrepancy between the predicted flows inside that ROI and the optimal, re-computed flows for that mask. We discard ROIs for which this discrepancy is large. Model ensembling is performed by averaging the flow predictions of 4 models trained separately. Image tiling and augmentation are performed together by dividing the image into overlapping tiles of the size of the training set patches (224 x 224 pixels). We use 50% overlaps for both horizontal and vertical tiling, resulting in every pixel being processed 4 times. We then combine the results by averaging the 4 flow predictions at every pixel, multiplied with appropriate taper masks to minimize edge effects. Furthermore, we take advantage of test data augmentation by flipping image tiles vertically, horizontally or in both directions depending on their position in the tile grid. The predicted flows are correspondingly flipped back with appropriately reversed sign before being averaged into the final predicted flows (see methods). An equivalent procedure is performed to augment the binary prediction of pixels inside/outside of cells.

#### Test time resizing

While it is easy to resize training set images to a common object size, such resizing cannot be directly performed on test images because we do not know the average size of objects in that image. In practice, a user might be able to quickly specify this value for their own dataset, but for benchmarks we needed an automated object size prediction algorithm. This information is not readily computable from raw images, but we hypothesized that the image style vectors might be a good representation from which to predict the object size. We predicted object sizes in two steps: 1) we train a linear regression model from the style vectors of the training set images, which is a good but not perfect prediction on the test set, 2) we refine the size prediction on the test set by running Cellpose segmentation after resizing to the object size predicted by the style vectors. Since this segmentation is already relatively good, we can use its mean object size as a better predictor of the true object size. We found that this refined object size prediction reached a correlation of 0.93 and 0.97 with the ground truth on test images, for the cytoplasm and nuclear dataset respectively (Figure S2). We include this algorithm as a calibration procedure which the user can choose to do either on every image, or just once for their entire dataset. We use this object size to resize test images, run the algorithm to produce the three output maps, and then resize these maps back to the original image sizes before running the pixel dynamics and mask segmentation. For all resizing operations we used standard bilinear interpolation from the OpenCV package [35].

#### Tiling and test augmentations

Image tiling is performed for three reasons: 1) to run Cellpose on images of arbitrary sizes, 2) to run Cellpose on the same image patch size used during training (224 by 224), and 3) to enable augmentation of each image tile in parallel with different augmentations on the other tiles. We start with an image of size *l_y_* by *l_x_* and divide this into 4 sets of tiles, where each tile set covers the entire image without overlap, except for some overlap at the right and bottom edges of the image. Each image tile of size 224 x 224 pixels can be completely specified by its upper-left corner position, defined by the coordinates (*r_y_* + 224 · *n_y_, r_x_* + 224 · *n_x_*), with *n_x_* = 0,1,2,.. and *n_y_* = 0,1,2,…, and (*r_x_,r_y_*) = (0,0), (0,112), (112,0), (112,112) respectively for the four sets of tiles. For the tiles at the edges which would contain pixels outside of cells, we move the entire tile in so it aligns with that edge of the image. For these 4 sets of tiles, we extract the corresponding image patches, and apply the augmentation transformation: we keep tile set 1 the same, we flip tile set 2 horizontally, we flip tile set 3 vertically, and we flip tile set 4 both horizontally and vertically. We then run Cellpose on all tiles, producing three output maps for each. The output maps are correspondingly flipped back, taking care to reverse the gradient directions for gradients along the flipped dimension. For example, in tile set 2 we flip horizontally both the horizontal and vertical gradients as well as the pixel probabilities, but we only reverse the sign of the horizontal gradients. We then use the tiles of the three outputs maps to reconstitute the gradient and probability maps for the entire image, tapering each tile around its 4 edges with a sigmoid taper with bandwidth parameter of 7.5 pixels. Note that in the final re-assembled image, almost all pixels are averages of 4 outputs, one for each tile set.

#### Mask quality threshold

Note there is nothing keeping the neural network from predicting horizontal and vertical flows that do not correspond to any real shapes at all. In practice, most predicted flows are consistent with real shapes, because the network was only trained on image flows that are consistent with real shapes, but sometimes when the network is uncertain it may output inconsistent flows. To check that the recovered shapes after the flow dynamics step are consistent with real masks, we recompute the flow gradients for these putative predicted masks, and compare them to the ones predicted by the network. When there is a large discrepancy between the two sets of flows, as measured by their mean squared difference, we exclude that mask since it is inconsistent. We cross-validate the threshold for this operation on a validation set held out from the main training dataset, and found that a value of 0.4 was optimal.

### 3D model

We extend the 2D model to 3D without the need for 3D labelled data by running the network on various slices of the 3D stack. Slices in XY provide X and Y flow information, slices in XZ provide X and Z flow information, and slices in YZ provide Y and Z flow information (Figure 5a). We average over the two estimates of each flow at each pixel (Figure 5d). From each slice we also predict the cell probability, and we average across these three estimates for each pixel. We threshold the cell probability at 0.5 and multiply it by the flows. We then use these flows to run the dynamics to create the mask estimates (Figure 5e). Objects smaller than 10% of the median cell volume (2000 voxels) were discarded (Figure 5f,g). The default median diameter was used (30 pixels).

### Benchmarking

We compared the segmentations performed by Cellpose to segmentations performed by Stardist [17] and Mask-RCNN [15, 16].

#### Training Stardist and Mask-RCNN

Stardist and Mask-RCNN were trained on the same training sets as Cellpose for the “specialized”, “generalized” and “nuclei” data sets. 12% of the training set was used for validation for each algorithm. Stardist and Mask-RCNN were trained for 1000 epochs. The learning rates were optimized to reduce the training error – this resulted in a learning rate of 0.0007 for Stardist and a learning rate of 0.001 for Mask-RCNN. For Mask-RCNN, TRAIN_ROIS_PER_IMAGE was increased to 300, MAX_GT_INSTANCES to 200, and DETECTION MAX INSTANCES to 400. Mask-RCNN was initialized with the pretrained “imagenet” weights. All other parameters and learning schedules for both algorithms were kept to their default values.

#### Using ilastik for 3D segmentation

As a comparison to Cellpose for 3D segmentation, we used ilastik [8] in a two-step process. First, each voxel was classified as ‘background’, ‘nucleus’, or ‘cytoplasm’ using a supervised algorithm. Then individual objects (i.e. cells) were identified using the hysteresis thresholding method. For the supervised classification, a small number of voxels were annotated manually and the following features were used for the classification: Gaussian smoothing (sigma = 0,3, 1,3.5 and 10), Gaussian gradient magnitude (sigma = 0.7, 1.6, and 5), difference of Gaussians (sigma = 0.7, 1.6, and 5), structure tensor eigenvalues (sigma = 0.7, 1.6, and 5), and Hessian of Gaussian eigenvalues (sigma = 0.7, 1.6, and 5). For the object identification, the ‘nucleus’ and ‘cytoplasm’ probability maps were used for setting the high (i.e. ‘core’) and low (ie. ‘final’) thresholds, respectively. Prior to thresholding, an isotropic Gaussian blur was applied (sigma = 1) and thresholds of 0.85 and 0.4 were chosen. Finally, objects smaller than 10% of the median cell volume (2000 voxels) were discarded.

#### Quantification of precision

We quantified the predictions of the algorithms by matching each predicted mask to the ground-truth mask that is most similar, as defined by the intersection over union metric (IoU). Then we evaluated the predictions at various levels of IoU; at a lower IoU, fewer pixels in a predicted mask have to match a corresponding ground-truth mask for a match to be considered valid. The valid matches define the true positives *TP*, the masks with no valid matches are false positives FP, and the ground-truth masks which have no valid match are false negatives *FN*. Using these values, we computed the standard average precision metric (AP) for each image:

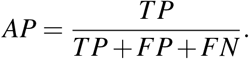

The average precision reported is averaged over the *AP* for each image in the test set, using the “match-ingjdataset” function from the Stardist package with the “by_image” option set to True [38].

---

**S1:**
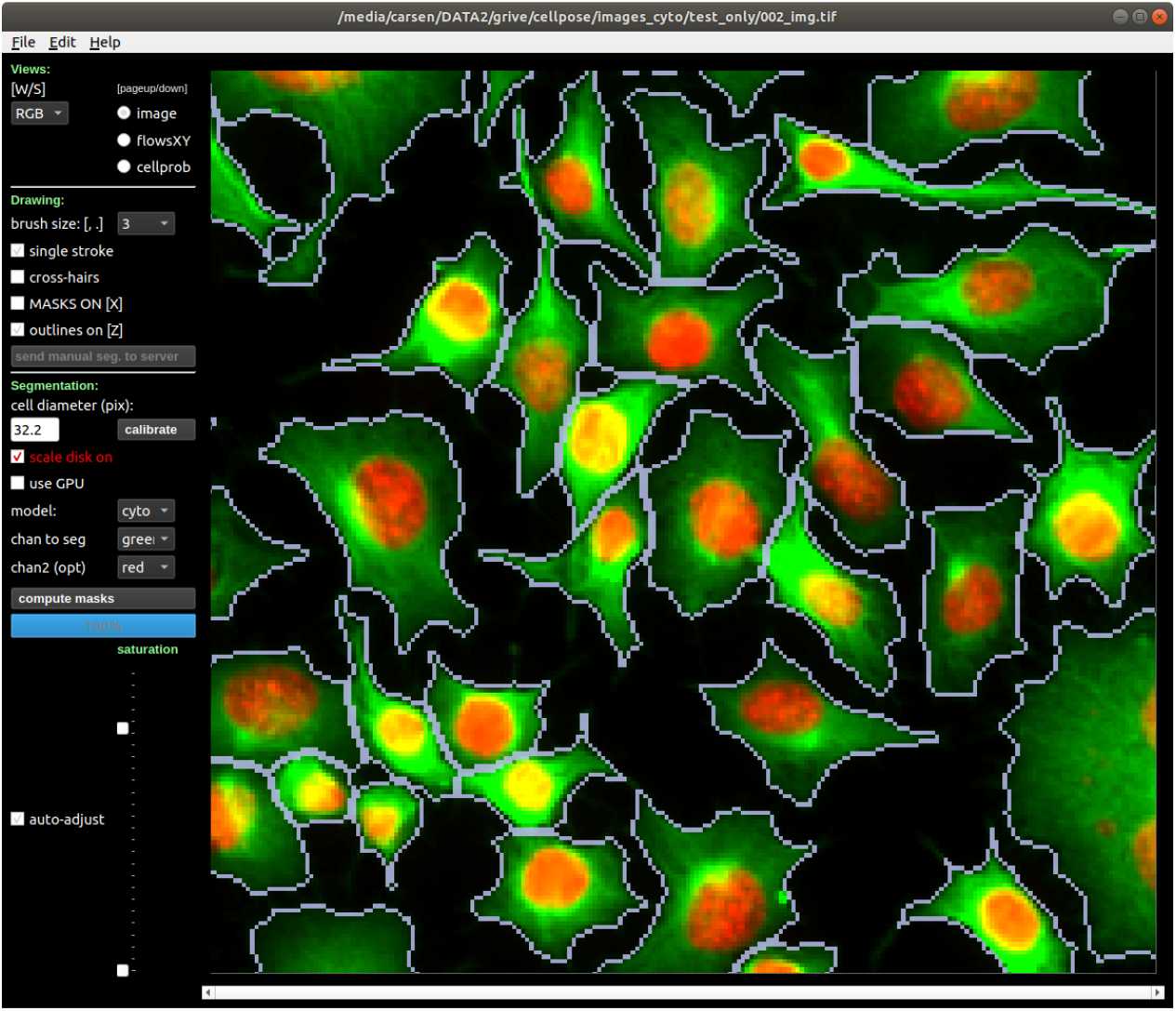
Graphical User Interface (GUI). Shown in the GUI is an image from the test set, zoomed in on an area of interest, and segmented using Cellpose. The GUI serves two main purposes: 1) easily run Cellpose “out-of-the-box” on new images and visualize the results in an interactive mode; 2) manually segment new images, to provide training data for Cellpose. The image view can be changed between image channels, cellpose vector flows and cellpose predicted pixel probabilities. Similarly, the mask overlays can be changed between outlines, transparent masks or both. The size calibration procedure can be run to estimate cell size, or the user can directly input the cell diameter, with an image overlay of a red disk shown as an aid for visual calibration. Dense, complete manual segmentations can be uploaded to our server with one button press, and the latest trained model can be downloaded from the dropdown menu.

**S2:**
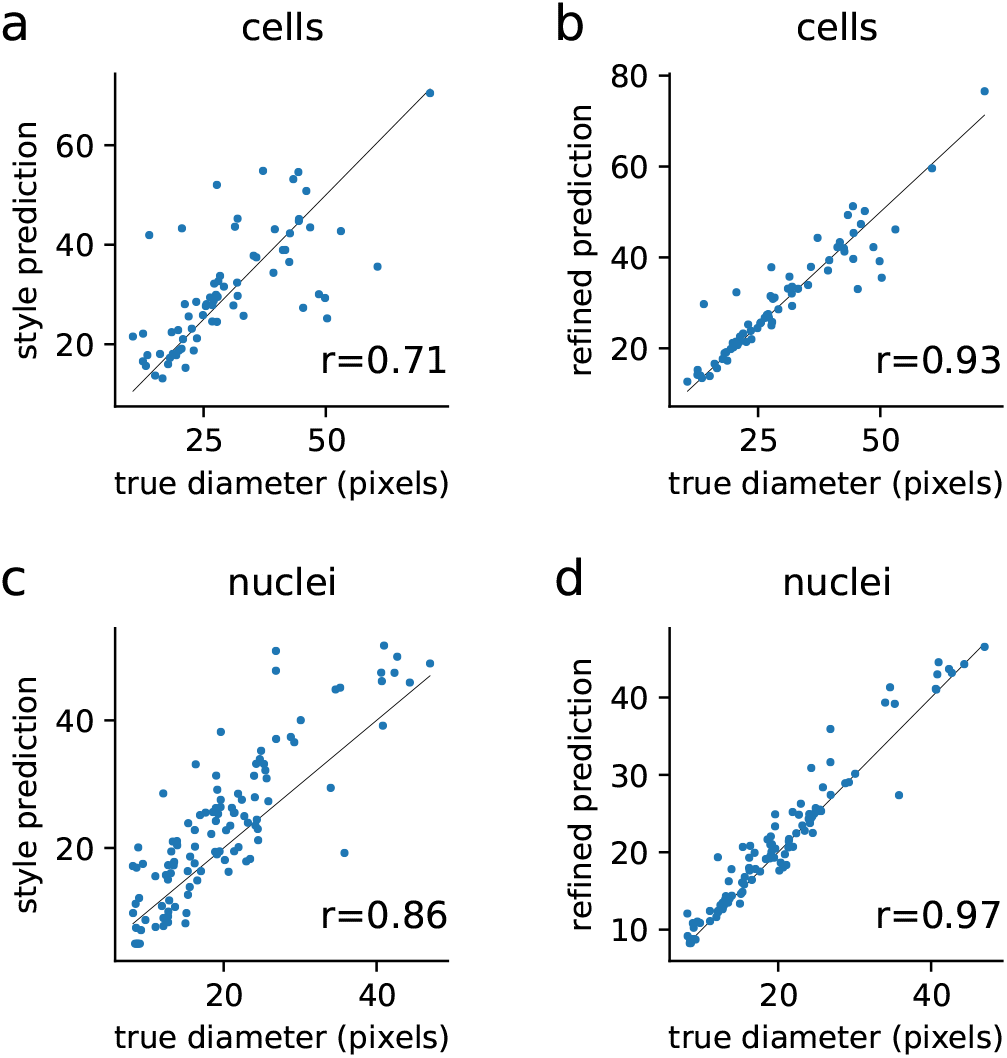
Prediction of median object diameter. **a**, The style vectors were used as linear regressors to predict the diameter of the objects in each image. Shown are the predictions for 69 test images, which were not used either for training cellpose or for training the size model. **b**, The refined size predictions are obtained after running cellpose on images resized according to the sizes computed in **a**. The median diameter of resulting objects is used as the refined size prediction for the final cellpose run. **cd,** Same as **ab** for the nucleus dataset.

**S3:**
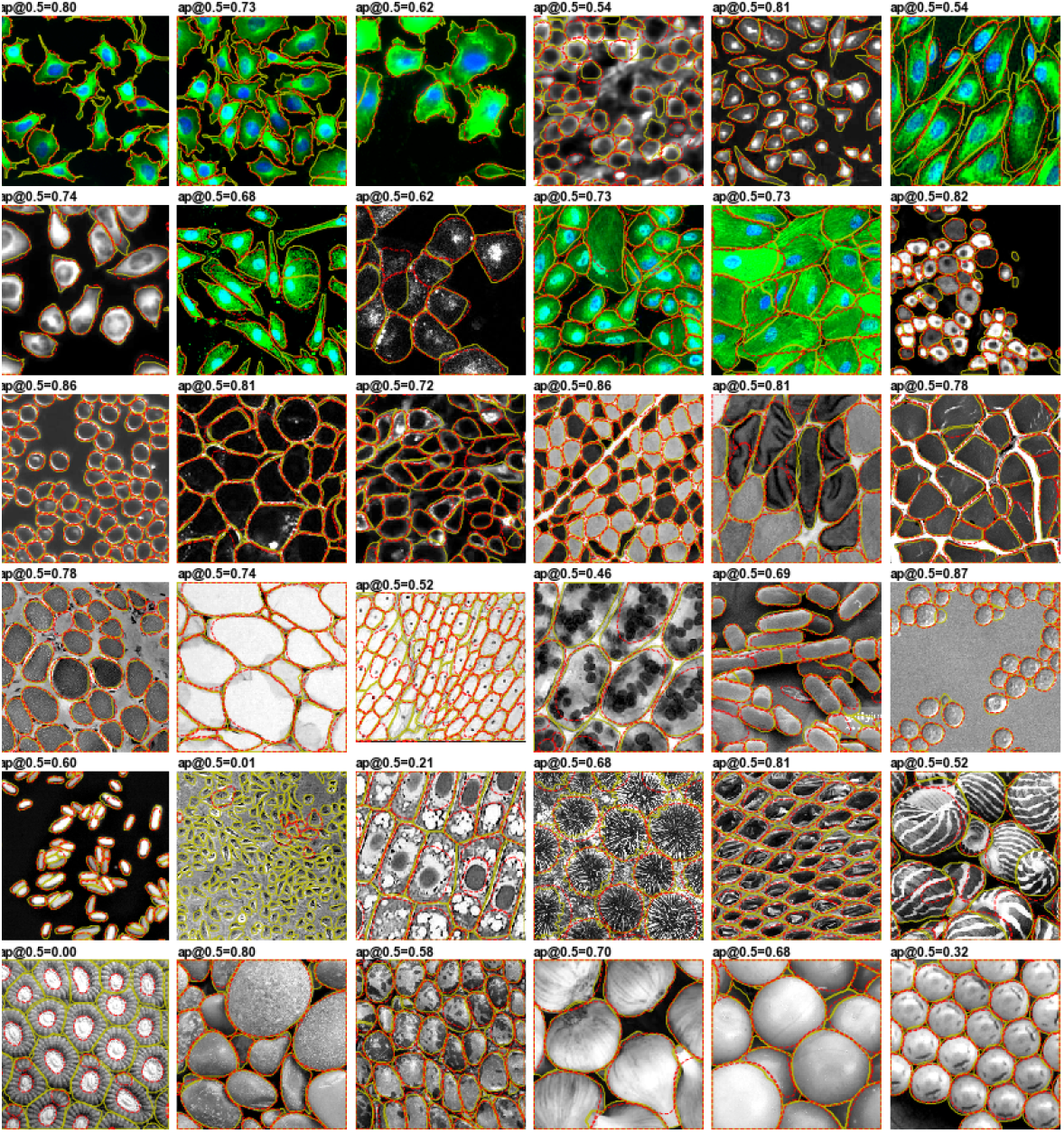
Example Stardist segmentations. Compare to Figure 4.

**S4:**
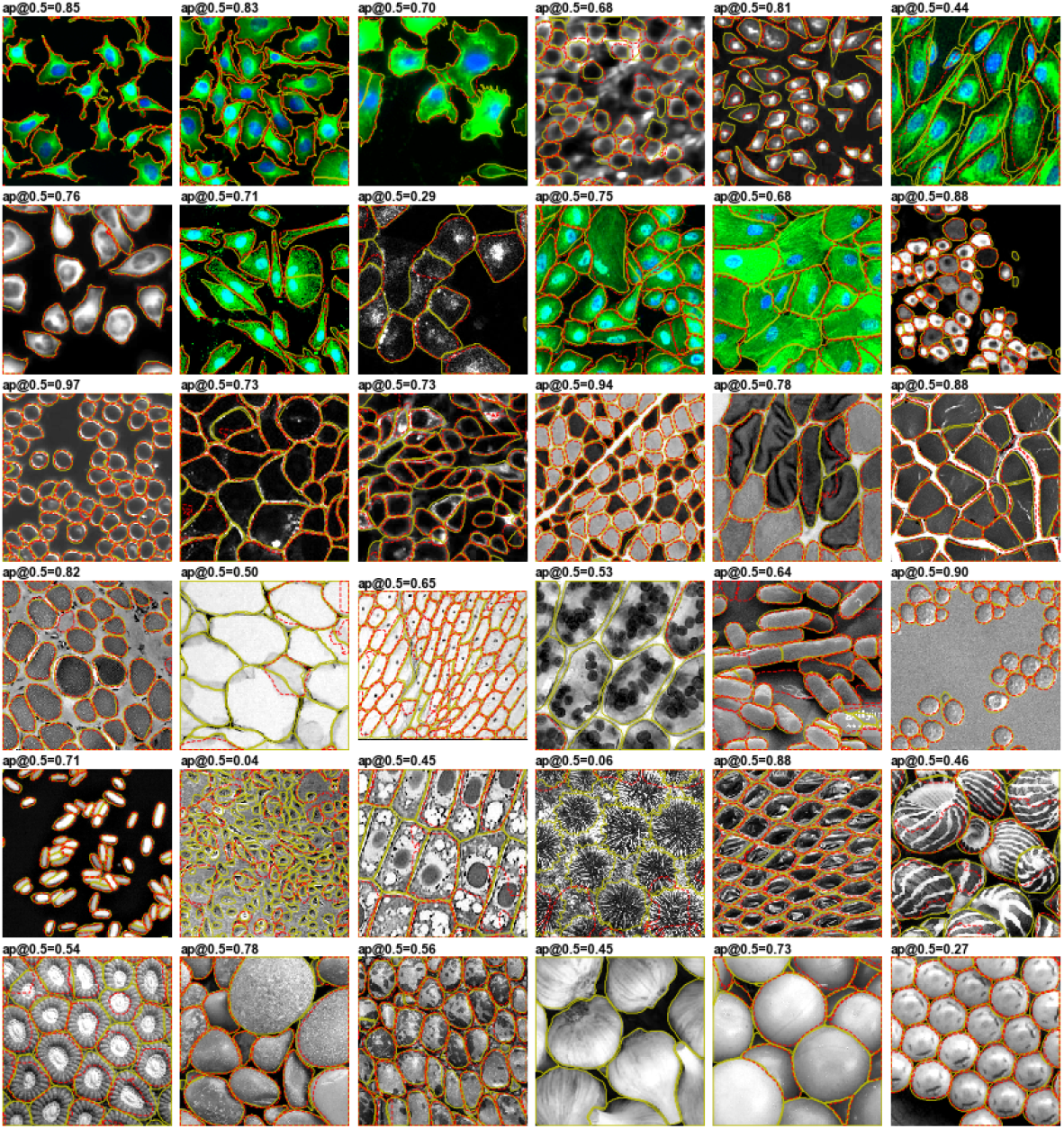
Example Mask R-CNN segmentations. Compare to Figure 4.

**S5:**
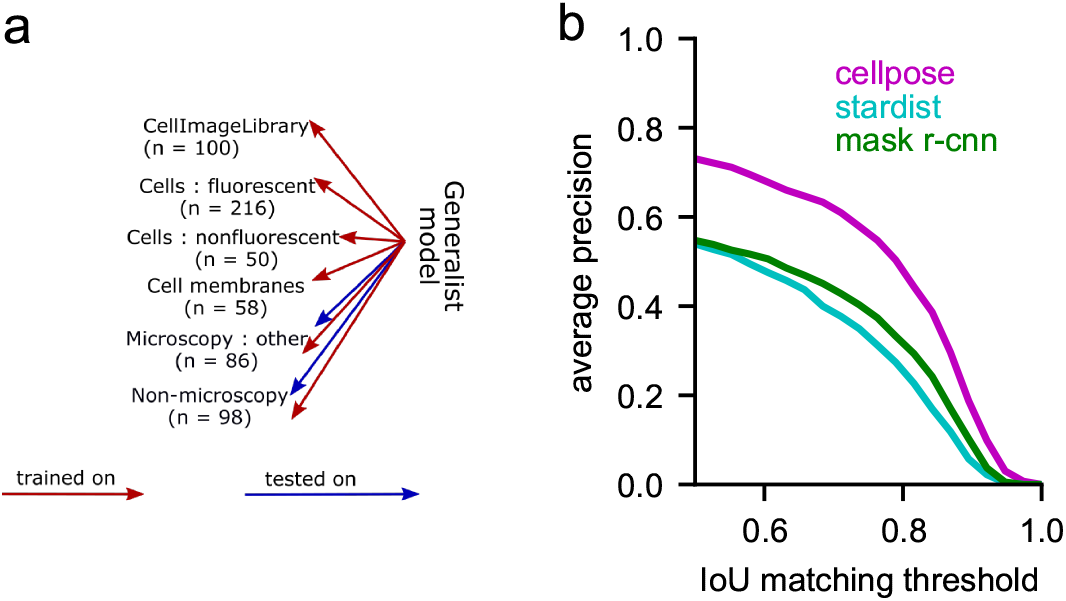
Performance on non-cell categories. **a**, We used the same trained models as shown in figure 3, and tested them on the more visually distinctive categories of “microscopy: other” and “non-microscopy”. Performance in this task can be an indicator of generalization performance, because most test images had no similar equivalent in the training dataset. **b**, Cellpose outperformed the other approaches by an even larger margin on this task, maintaining accuracy scores in the range of 0.7-0.8 at an IoU threshold of 0.5. These scores show that Cellpose can be useful as an all-purpose segmentation algorithm for groups of similar objects.

**S6:**
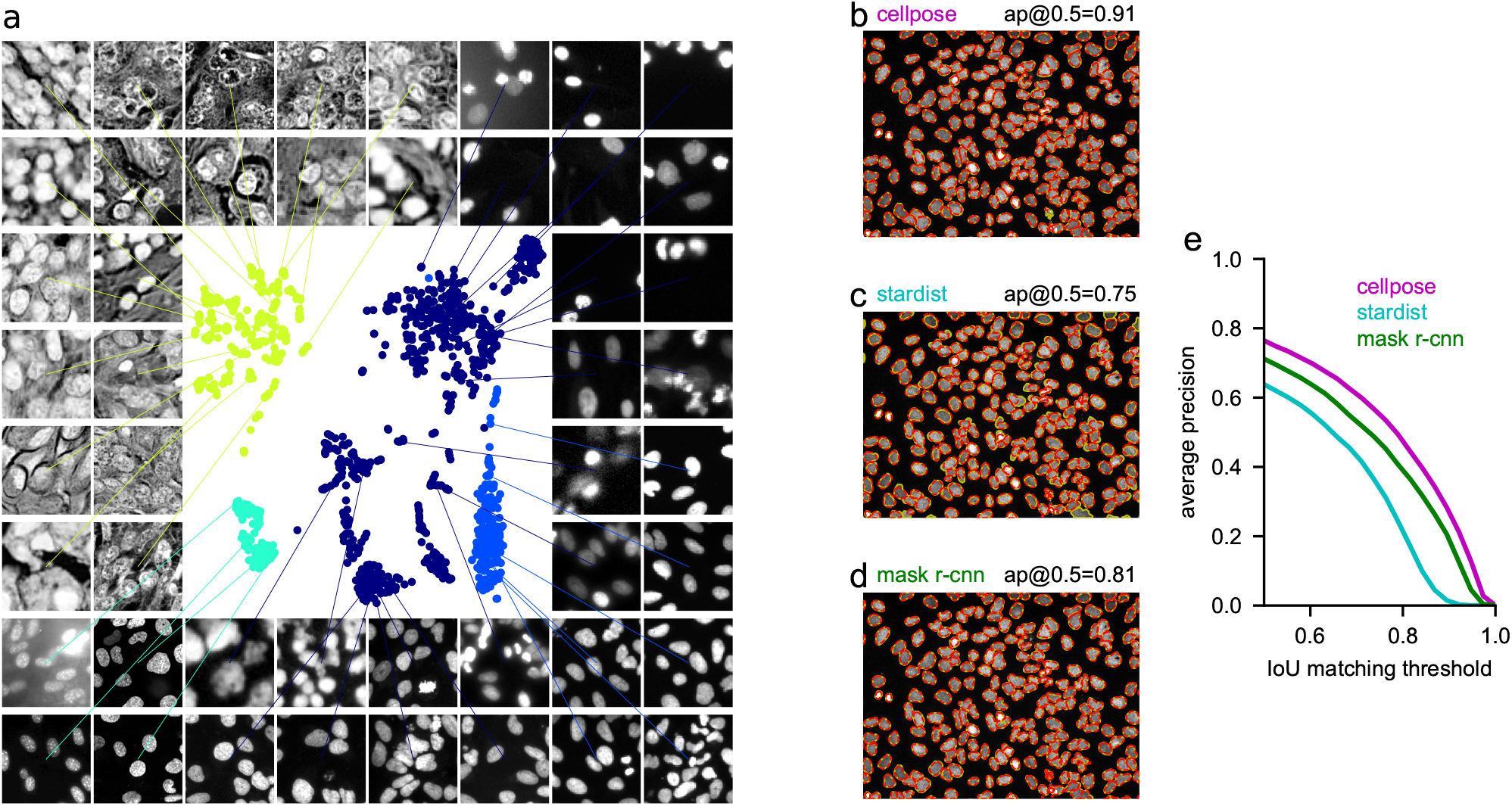
Benchmarks for dataset of nuclei. **a,** Embedding of image styles for the nuclear dataset of 1139 images, with a new Cellpose model trained on this dataset. Legend: dark blue = Data Science Bowl dataset, blue = extra kaggle, cyan = ISBI 2009, green = MoNuSeg. **bcd,** Segmentations on one example test image. **e,** Accuracy scores on test data.

**S7:**
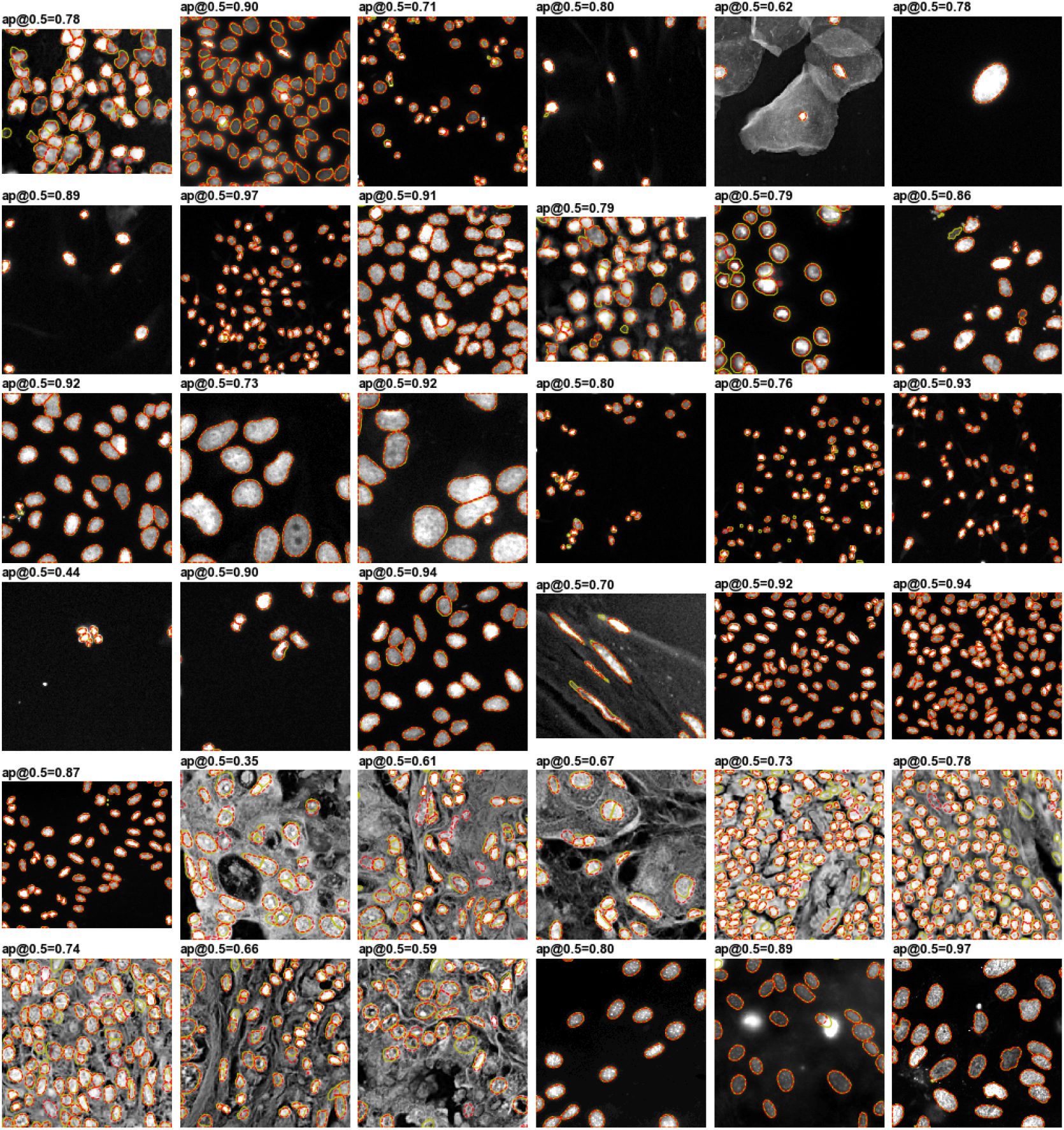
Example cellpose segmentations for nuclei. The ground truth masks segmented by a human operator are shown in yellow, while the predicted masks are shown in dotted red line.

