## Supplementary figures and images for "Cellpose: a generalist algorithm for cellular segmentation"

### Movie 1

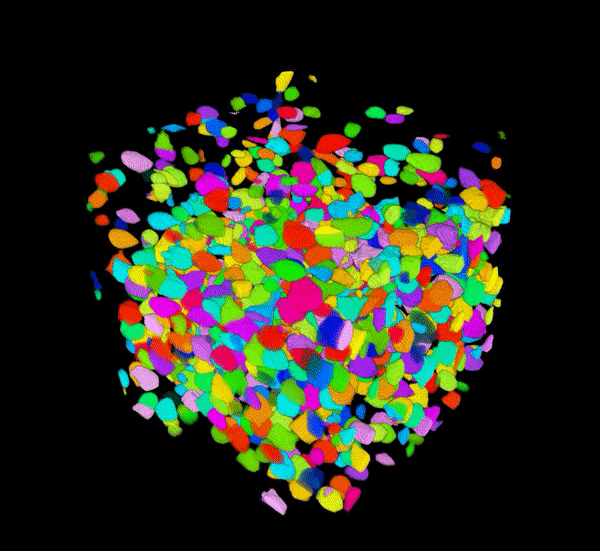

### Movie 2

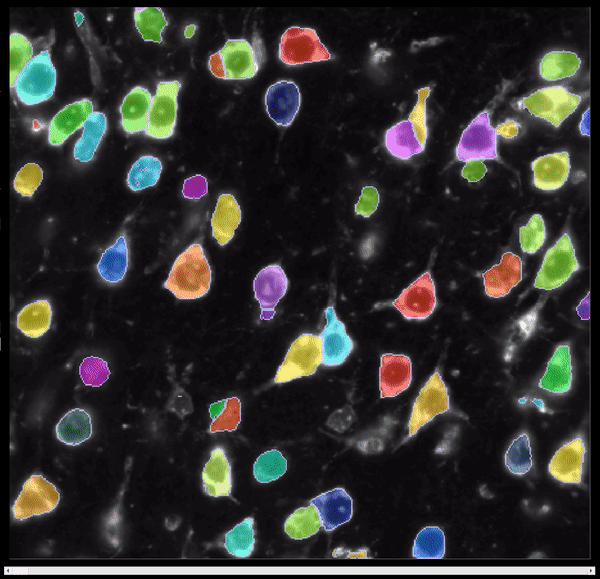
